# DETIRE: A Hybrid Deep Learning Model for identifying Viral Sequences from Metagenomes

**DOI:** 10.1101/2021.11.19.469211

**Authors:** Yan Miao, Fu Liu, Tao Hou, Qiaoliang Liu, Tian Dong, Yun Liu

**Affiliations:** College of Communication Engineering, Jilin University, Changchun, Jilin Province, China

## Abstract

A metagenome contains all DNA sequences from an environmental sample, including viruses, bacteria, fungi, actinomycetes and so on. Since viruses are of huge abundance and have caused vast mortality and morbidity to human society in history as a kind of major pathogens, detecting viruses from metagenomes plays a crucial role in analysing the viral component of samples and is the very first step for clinical diagnosis. However, detecting viral fragments directly from the metagenomes is still a tough issue because of the existence of huge number of short sequences. In this paper, a hybrid Deep lEarning model for idenTifying vIral sequences fRom mEtagenomes (DETIRE), is proposed to solve the problem. Firstly, the graph-based nucleotide sequence embedding strategy is utilized to enrich the expression of DNA sequences by training an embedding matrix. Then the spatial and sequential features are extracted by trained CNN and BiLSTM networks respectively to improve the feature expression of short sequences. Finally, the two set of features are weighted combined for the final decision. Trained by 220,000 sequences of 500bp subsampled from the Virus and Host RefSeq genomes, DETIRE identifies more short viral sequences (<1,000bp) than three latest methods, DeepVirFinder, PPR-Meta and CHEER. DETIRE is freely available at https://github.com/crazyinter/DETIRE.

## Introduction

High-Throughput Sequencing or called Next-Generation Sequencing (NGS) technology, which makes it possible to obtain the whole nucleotide sequences directly from environmental samples, has played important roles in many fields, such as pathogen detection [1, 2] and human disease analysis [3–5]. In these applications, detecting viruses from metagenomic sequences becomes more and more essential because it is the first step in the analysis of viruses [6]. However, it is still a quite difficult task because of their relatively low abundances compared to those of bacteria and high mutation rates.

To overcome this challenge, several methods have been being proposed to identify viruses from metagenomes in the past several years, and can be categorized into similarity-based, machine learning-based and deep learning-based methods. Similarity-based methods generally map a query sequence to a reference data set and recognize it as the one with the highest similarity score [7–17]. These methods, however, suffer from long execution time during the mapping process, and hardly detect short viral sequences because of the limited features they have. Different to the similarity-based methods, machine learning-based methods could extract human-designed features from DNA sequences and classify them by a well-trained classifier, such as VirFinder [18], MARVEL [19] and VirSorter2 [20]. Although these methods have the ability to produce more accurate results, the features have to be designed artificially and the performance on identifying short viral sequences is relatively poor.

With the great success of deep learning methods in the past few years, several deep learning-based methods have been proposed to identify viruses from metagenomes. Long-short term memory (LSTM) network and convolutional neural network (CNN) are the most commonly used models. For example, ViraMiner [21], VirNet [22] and RNN-VirSeeker [23] utilize a single LSTM network to learn the interconnections between each part in a one-hot encoded sequence; DeepVirFinder [24], PPR-Meta [25] and CHEER [26] establish a single CNN to extract high-level features from one-hot encoded sequences automatically before a set of dense layers and a softmax layer for classification. However, the following two issues hamper the performance of deep learning models for the recognition of short viral sequences: 1) the single deep learning architecture suffers from failing to extract enough features to represent sequences; 2) the one-hot encoding strategy omits the relationship between two parts of a sequence because of its orthogonal property [27].

To solve the above-mentioned issues, a novel hybrid deep learning-based virus identifier, namely DETIRE, is proposed in this paper to identify viral fragments directly from metagenomes. DETIRE is a two-stage architecture, containing a graph convolutional network (GCN) based sequence embedder and a two-path deep learning model. First, every sequence is cut into several 3-mer fragments, which are then successively input to the GCN-based sequence embedder to train the representations of all 3-mer fragments. After that, these embedded fragments are then fed into the CNN-path and BiLSTM-path to learn their features, respectively. Finally, by two dense layers and a softmax layer, a pair scores are generated, and the higher score determines which type the input sequence is.

The main contributions of this paper are two folds: 1) the GCN-based embedder is utilized to enrich the representations of short sequences; 2) BiLSTM and CNN are combined to learn not only the spatial characteristics but also sequential characteristics simultaneously to generate abundant features of short sequences. To the best of our knowledge, DETIRE is the first hybrid model that combines CNN and BiLSTM networks to identify viruses from metagenomes.

The last of this paper is organised as follows: Section **Materials and methods** introduces the datasets used to train and test the DETIRE, and details the architecture of DETIRE and its training strategy. Section **Results and Discussion** show the performance of the DETIRE on several datasets and discusses the selection of some key parameters which can affect the overall performance. Section **Conclusions** gives a brief conclusion of this paper.

## 1 Materials and Methods

### 1.1 Virus and host RefSeq genome datasets for training and testing

Virus RefSeq genome (up to October 18, 2020) was downloaded from NCBI Virus (https://www.ncbi.nlm.nih.gov/labs/virus/vssi/#/virus?SeqType_s=Genome&SourceDB_s=RefSeq). It has been proved that the less difference between the lengths of sequences from training and testing set, the better classification result will be achieved [19]. Thus all 13,274 viral sequences were split into a set of non-overlapped fragments with a length of 500bp, resulting 778,390 fragments totally. The whole set of 500bp viral fragments combined with 770,000 sequences of 500bp subsampled from 4,410 prokaryotic host RefSeq genomes supplied in VirFinder [19], calling the GCN-training dataset, were jointly used to train the GCN-based model. Then 110,000 viral sequences and 110,000 host sequences were randomly subsampled from the fragments above. Then they were divided into the training and testing datasets at a scale of 10:1. The NCBI accession numbers of the viral RefSeqs can be found at https://github.com/crazyinter/DETIRE/blob/main/supplementaryfiles/Virus_RefSeq_accession_numbers.csv.

### 1.2 Composition of DETIRE

DETIRE utilizes a two-stage strategy for virus prediction, including GCN-based sequence embedding and deep learning-based sequence classification (Fig 1). Before embedding, every 3-mer fragment is generated by a three-bases sliding window moving from the head to the tail of the sequence with a stride of one. For example, the original nucleotide sequence ‘ATTGCCTGACAT’ will be cut into ‘ATT, TTG, TGC, GCC, CCT, CTG, TGA, GAC, ACA, CAT’.

**Fig 1.**
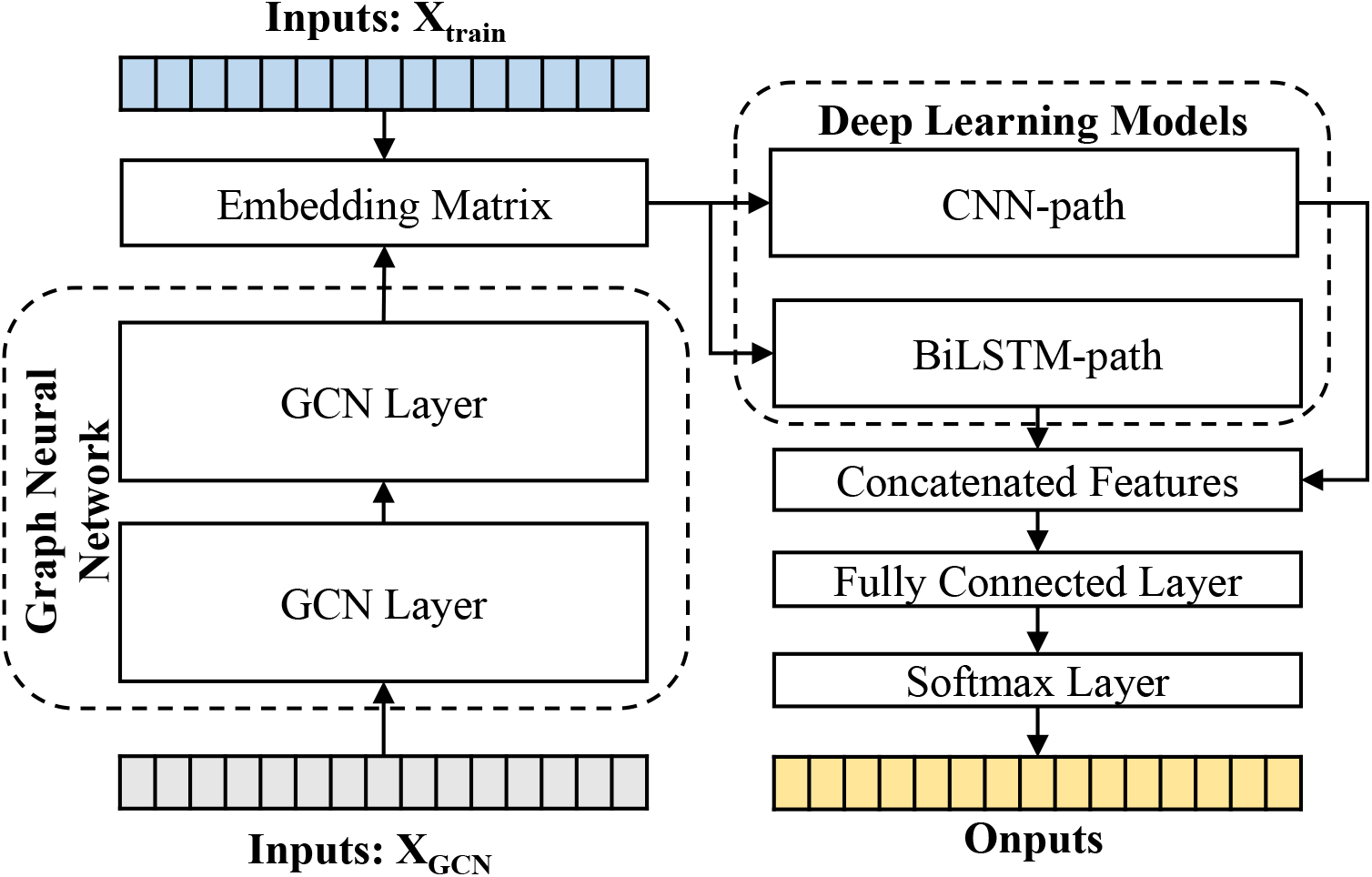
The workflow of DETIRE. DETIRE contains GCN based sequence embedding model and a deep learning-based method to learn features automatically of viral sequences and identify them directly from metagenomes. First, the graph neural network learns the high-level representations of 3-mer fragments in each sequence through supervised back propagation. Then DETIRE extracts the features of spatial characteristics and sequential characteristics by designed CNN model and LSTM model, respectively. Finally, the learned features are combined together to make the final decision by several dense layers and a softmax layer.

In the process of sequence embedding, TextGCN [28] is utilized to learn the meaningful high-level representations of all 3-mer fragments from every nucleotide sequence. Firstly, a heterogeneous graph containing 3-mer-fragment nodes is built in order to model global co-occurrence between these 3-mer fragments explicitly. Then the built graph is fed into a simple two-layer GCN [29]. The first layer constructs the nodes and edges. Every nucleotide sequence in the GCN-training dataset and all unique 3-mer fragments from it are constructed to their single nodes. There are no edges between each nucleotide sequence. Edges are built between 3-mer fragments and their original sequences. All 64 3-mer fragments have an edge between each of them (Fig 2). The weight of the edge between a sequence node and a 3-mer fragment node is determined by the term frequency-inverse document frequency (TF-IDF) [30] of the fragment in the sequence, where term frequency is the frequency of the 3-mer fragment appears in the sequence and inverse document frequency is the logarithmically scaled inverse fraction of the number of sequences that contain the 3-mer fragment. Point-wise mutual information (PMI) [31], a popular measure for word associations, is employed to calculate weights between two fragment nodes. The second layer learns the fragment and sequence embeddings in each node. Finally, these nodes are fed into a softmax classifier, after which the cross-entropy error over all labelled sequences is defined as the cost function [32]. After 500 epochs of backpropagation by the Adam optimization algorithm [33] with a learning rate of 0.022 and a dropout rate of 0.5, the 30-dimension representations of all 3-mer fragments in the second layer of the GCN are embedded into the sequences in the training and testing datasets.

**Fig 2.**
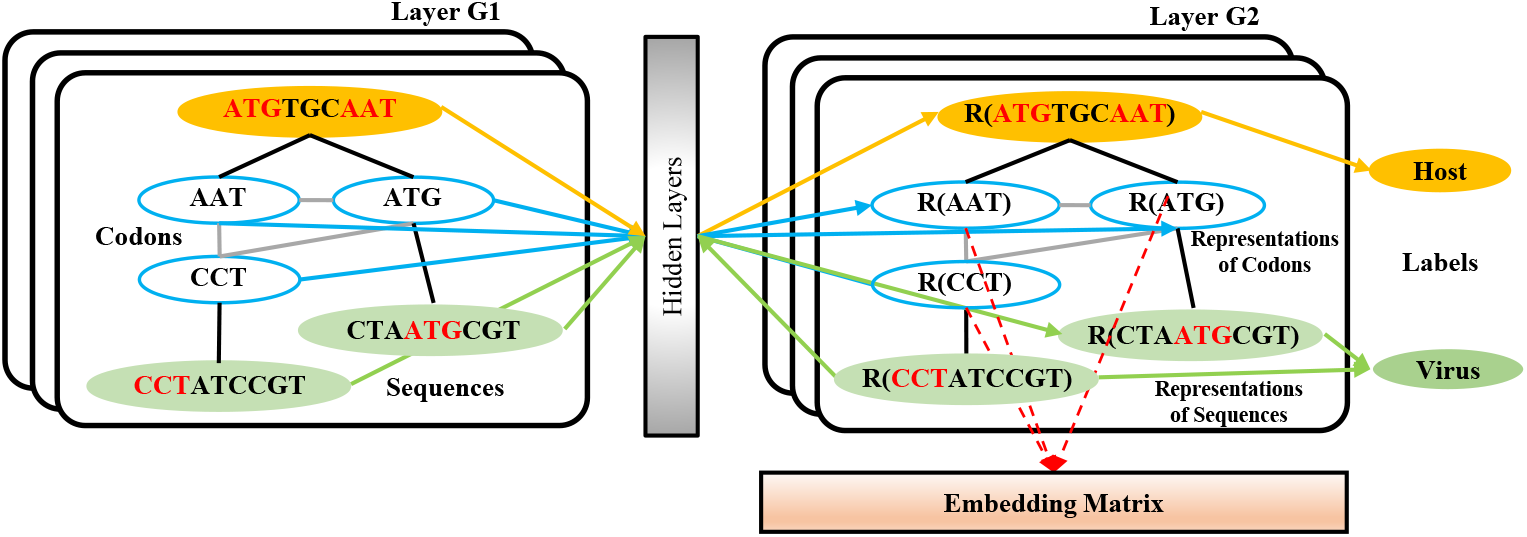
The structure of the GCN-based embedding model. Every nucleotide sequence in the GCN-training dataset and all unique 3-mer fragments from it are constructed to their single nodes. There are no edges between each nucleotide sequence. Edges are built between 3-mer fragments and their original sequences. All 64 3-mer fragments have an edge between each of them. After training strategy, the vectors from the codon nodes in the second GCN layer are combined to a embedding matrix.

In the process of sequence classification, two parallel deep learning models, CNN and BiLSTM, are respectively used to learn spatial and sequential features of sequences. In the CNN path, each embedded sequence is considered as an image to extract a spatial feature through three sets of layers. Each set of the layer contains a convolutional layer (16, 32 and 64 filters with size of 7*7, 5*5 and 3*3, respectively), a ReLU [34] activation function, a max pooling layer (with a pooling size of 4*4 and a stride of 4), and a batch normalization (BN) layer, respectively. In the BiLSTM path, the embedded 3-mer fragments in a sequence are input into the BiLSTM cells (498 tokens totally) one by one, generating a sequential feature. Then the first dense layer with 100 hidden neurons receives the weighted merged two sets of features from the CNN path and BiLSTM path. The second hidden layer after that contains 30 hidden neurons. Finally, a softmax layer generates two scores that reflect the likelihood of the input sequence as a virus or not. The weights of merging are two sets of trainable parameters which can be finetuned during the training progress. All of the parameters here are updated by Adam [33] optimizer with a mini-batch of 200 for 20 epochs to reduce the cross-entropy loss with a learning rate of 0.03.

### 1.3 Evaluation criteria

Universally, a confusion matrix is calculated to evaluate the performance of a classifier according to four statistics: True Positives (TP), False Positives (FP), True Negatives (TN), and False Negatives (FN) [23]. TP are examples correctly labeled as positives; FP refer to negative examples incorrectly labeled as positive; TN correspond to negatives correctly labeled as negative; and FN refer to positive examples incorrectly labeled as negative. Several high-level criteria are further calculated based on the confusion matrix, such as recall, accuracy, precision and F1 score:

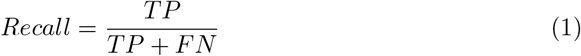

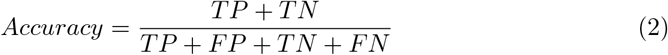

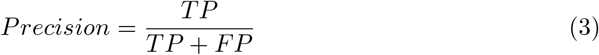

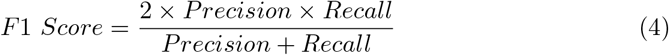

## 2 Results and Discussion

### 2.1 A marine metagenome dataset from CAMI

To test the performance of DETIRE on identifying viral sequences, a marine metagenome was downloaded from the 2nd CAMI Challenge Marine Dataset (https://data.cami-challenge.org/participate), a short and long read shotgun metagenome data from samples at different seafloor locations of a marine environment. All sequences from the fasta document named CAMI2_short_read_pooled_gold_standard_assembly were selected. These sequences were then mapped to the Virus RefSeq genome using BLAST (Basic Local Alignment Search Tool) [35] with default parameters. As a result, 6,262 sequences shorter than 500bp, 5,000 sequences in length of 500-1,000bp, 6,636 sequence in length of 1,000-3,000bp, and 9,365 sequences longer than 3,000bp were recognized as viral sequences. The same amount of non-viral short sequences was also randomly subsampled from the rest sequences for each length, respectively. Together, these 54,526 mixed sequences were built into a metagenome dataset to test DETIRE.

### 2.2 A real human gut metagenome dataset

A real human gut metagenome dataset was downloaded from the NCBI short-read archive (accession ID: SRA052203) [36]. The same BLAST progress was done as it was in the 2nd CAMI Challenge Marine Dataset, resulting 301 viral sequences shorter than 500bp, 343 sequences in length of 500-1,000bp, 639 sequence in length of 1,000-3,000bp, and 865 sequences longer than 3,000bp. Similarly, the same amount of non-viral short sequences for each length was subsampled randomly from the rest non-viral sequences. Totally, 4,296 mixed sequences were used to test DETIRE.

### 2.3 The influence of k-mer fragments and their embedding sizes on the viral identification results

Every nucleotide sequence was converted into a set of k-mer fragments before being expressed as high-level vectors by the graph neural network to enrich the representation of the sequence. The chosen of the k value and the embedding size have an significant impact on the identification performance. To find a relatively optimal set of the two parameters, a lot of models have been trained by the established training dataset and evaluated by the testing dataset. Each model corresponds to a specific set of parameters, where k is chosen from [1, 2, 3, 4, 5, 6, 7, 8, 9, 10] and embedding size is chosen from [4, 8, 30, 50, 100, 200, 500, 1,000], respectively. The evaluation results are shown in Fig 3. When the embedding size is a little lower than 4k (all combinations of the k-mer fragments), the AUROC value is close to the maximum. Thus 3-mer fragments with the embedding size of 30 was chosen into our final model. The final representations of all 3-mer fragments can be found at https://github.com/crazyinter/DETIRE/blob/main/supplementary%20files/embedding_maitrix.csv.

**Fig 3.**
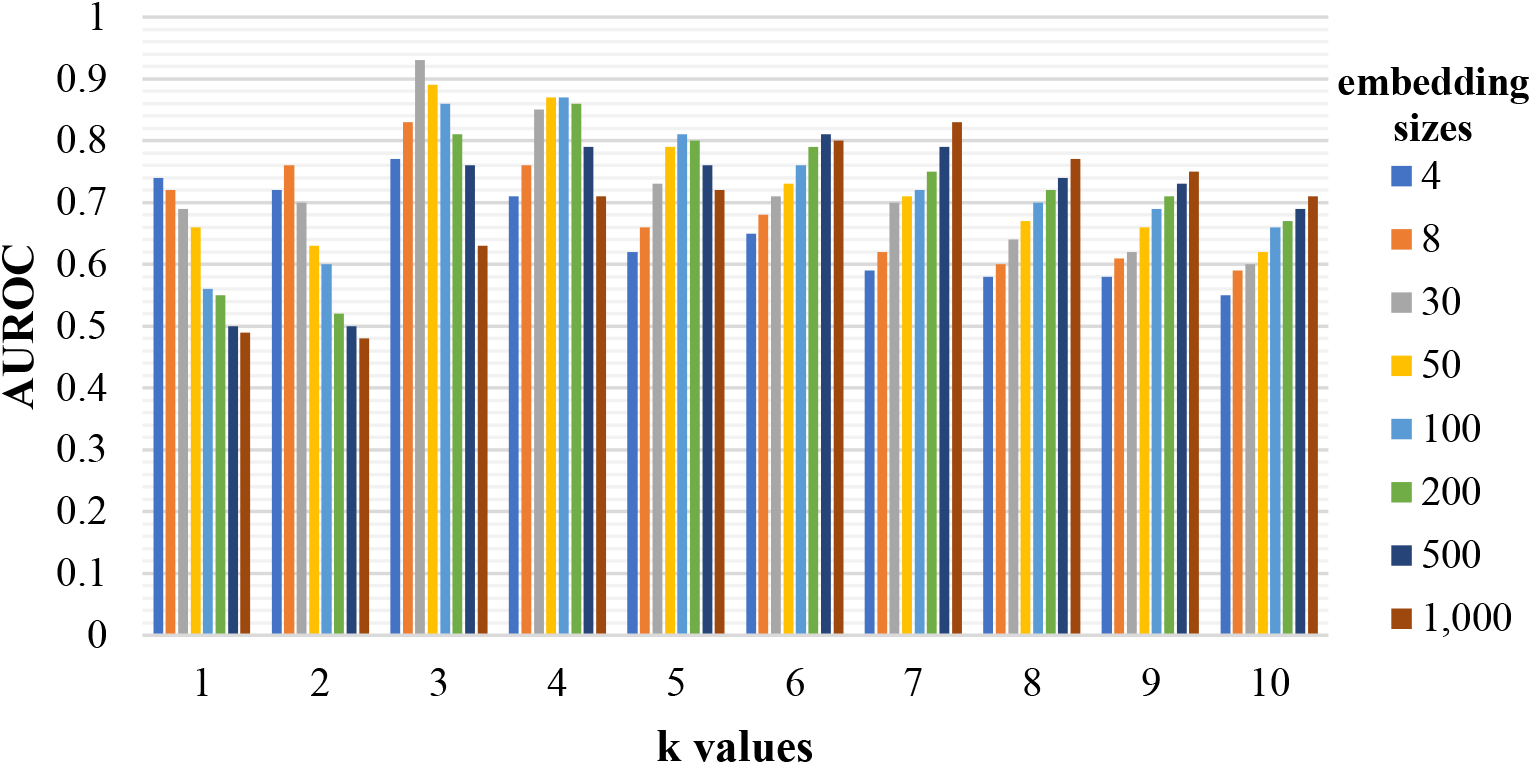
The performance of the models with various k values and embedding sizes on the testing dataset. Sequences with specific k-mer fragments and embedding sizes were used to train several models, which were then evaluated by the testing dataset. The AUROC values of these models were calculated. The k is chosen from [1, 2, 3, 4, 5, 6, 7, 8, 9, 10] and the embedding size is chosen from [4, 8, 30, 50, 100, 200, 500, 1,000].

### 2.4 Performance on the testing dataset

To prove the outstanding performance of DETIRE on identifying short viral sequences (<500bp), a testing experiment on the testing dataset was done to make a comparison between DETIRE and three latest viral identification methods, DeepVirFinder, PPR-Meta and CHEER. The accuracies, recalls, precisions and F1 scores of the four methods are calculated in Table 1. DETIRE outperforms DeepVirFinder and PPR-Meta at all of the four criteria. Although it achieves 0.0009 lower value of precision than CHEER which is caused by a little more misclassified non-viral sequences, DETIRE performs better at the other three criteria, 0.0008, 0.0034 and 0.0012 higher at accuracy, recall and F1 score, respectively.

**Table 1.** Comparison of Accuracies, Recalls, Precisions and F1 Scores of DeepVirFinder, PPR-Meta, CHEER and DETIRE on the testing dataset.

| Criteria | DeepVirFinder | PPR-Meta | CHEER | DETIRE |
| --- | --- | --- | --- | --- |
| Accuracy | 0.8297 | 0.8306 | 0.8464 | **0.8472** |
| Recall | 0.8292 | 0.8323 | 0.8472 | **0.8506** |
| Precision | 0.8299 | 0.8294 | **0.8458** | 0.8449 |
| F1 score | 0.8296 | 0.8308 | 0.8465 | **0.8477** |

Bold values represent the best performance.

**Table 2.** Comparison of Accuracies, Recalls, Precisions and F1 Scores of DeepVirFinder, PPR-Meta, CHEER and DETIRE on the CAMI Marine metagenome.

| Length | Criteria | Deep-VirFinder | PPR-Meta | CHEER | DETIRE |
| --- | --- | --- | --- | --- | --- |
| <500bp | Accuracy | 0.6823 | 0.7004 | 0.7327 | **0.7501** |
|  | Recall | 0.6808 | 0.7014 | 0.7340 | **0.7542** |
|  | Precision | 0.6828 | 0.7000 | 0.7321 | **0.7480** |
|  | F1 score | 0.6838 | 0.6995 | 0.7330 | **0.7511** |
| 500-1,000bp | Accuracy | 0.6947 | 0.7189 | 0.7503 | **0.7700** |
|  | Recall | 0.6954 | 0.7160 | 0.7482 | **0.7696** |
|  | Precision | 0.6944 | 0.7202 | 0.7514 | **0.7702** |
|  | F1 score | 0.6949 | 0.7181 | 0.7498 | **0.7699** |
| 1,000-3,000bp | Accuracy | 0.7161 | 0.7170 | **0.7395** | 0.7374 |
|  | Recall | 0.7105 | 0.7092 | **0.7416** | 0.7322 |
|  | Precision | 0.7185 | 0.7205 | 0.7384 | **0.7399** |
|  | F1 score | 0.7145 | 0.7148 | **0.7400** | 0.7360 |
| >3,000bp | Accuracy | 0.7251 | 0.7395 | **0.7442** | 0.7292 |
|  | Recall | 0.7225 | 0.7414 | **0.7461** | 0.7320 |
|  | Precision | 0.7263 | 0.7385 | **0.7433** | 0.7279 |
|  | F1 score | 0.7244 | 0.7400 | **0.7447** | 0.7299 |
| Overall | Accuracy | 0.7075 | 0.7213 | 0.7415 | **0.7435** |
|  | Recall | 0.7050 | 0.7197 | 0.7426 | **0.7440** |
|  | Precision | 0.7085 | 0.7219 | 0.7410 | **0.7432** |
|  | F1 score | 0.7068 | 0.7208 | 0.7418 | **0.7436** |

Bold values represent the best performance.

**Table 3.** Comparison of Accuracies, Recalls, Precisions and F1 Scores of DeepVirFinder, PPR-Meta, CHEER and DETIRE on the real human gut metagenome.

| Length | Criteria | DeepVirFinder | PPR-Meta | CHEER | DETIRE |
| --- | --- | --- | --- | --- | --- |
| <500bp | Accuracy | 0.7558 | 0.7226 | 0.8904 | **0.9203** |
|  | Recall | 0.7475 | 0.711 | 0.8837 | **0.907** |
|  | Precision | 0.7601 | 0.7279 | 0.8956 | **0.9317** |
|  | F1 score | 0.7538 | 0.7193 | 0.8896 | **0.9192** |
| 500-1,000bp | Accuracy | 0.7624 | 0.7711 | 0.8907 | **0.895** |
|  | Recall | 0.7609 | 0.7668 | 0.8892 | **0.9009** |
|  | Precision | 0.7632 | 0.7735 | **0.8918** | 0.8905 |
|  | F1 score | 0.762 | 0.7701 | 0.8905 | **0.8957** |
| 1,000-3,000bp | Accuracy | 0.795 | 0.8169 | **0.892** | 0.8701 |
|  | Recall | 0.7919 | 0.8138 | **0.8936** | 0.8795 |
|  | Precision | 0.7969 | 0.8189 | **0.8908** | 0.8633 |
|  | F1 score | 0.7981 | 0.8163 | **0.8922** | 0.8713 |
| >3,000bp | Accuracy | 0.7977 | 0.852 | **0.8988** | 0.852 |
|  | Recall | 0.7954 | 0.8566 | **0.8971** | 0.8659 |
|  | Precision | 0.7991 | 0.8488 | **0.9002** | 0.8425 |
|  | F1 score | 0.7972 | 0.8527 | **0.8987** | 0.854 |
| Overall | Accuracy | 0.7854 | 0.8105 | **0.8943** | 0.8648 |
|  | Recall | 0.7821 | 0.8091 | **0.8929** | 0.8813 |
|  | Precision | 0.7873 | 0.8114 | **0.8954** | 0.8531 |
|  | F1 score | 0.7847 | 0.8103 | **0.8957** | 0.867 |

Bold values represent the best performance.

**Table 4.** Comparison of the testing time consuming on the three datasets.

|  | Datasets | DeepVirFinder | PPR-Meta | CHEER | DETIRE |
| --- | --- | --- | --- | --- | --- |
| Testing time (s) | Testing dataset | 91 | 85 | 88 | **76** |
|  | CAMI Marine | 332 | 298 | 305 | **287** |
|  | Real human gut | 122 | 107 | 119 | **102** |

Bold values represent the best performance.

### 2.5 Performance on the CAMI Marine metagenome

To deal with different lengths of sequences from the metagenome and to avoid vanishing gradient problem in the process of training BiLSTM, a sequence longer than 500bp will be divided into several non-overlapped sub-sequences of 500bp before input into BiLSTM path. If the length of the last part in the sequence is shorter than 500bp, the last bases of the sequence will be zero-padded and be regarded as a single subsequence. Then all of the sub-sequences are input to the hybrid deep learning model one after another to get their own scores, the average of which will be the final score and is contributed to identifying whether the query long sequence is viral or not.

The accuracies, recalls, precisions and F1 scores of DETIRE, CHEER, PPR-Meta, and DeepVirFinder on classifying viral and non-viral sequences from the CAMI Marine metagenome are calculated and made a comparison in Table **??**. DETIRE exceeds DeepVirFinder at all of the four criteria on identifying all lengths of viral sequences. DETIRE is better than PPR-Meta when the length is shorter than 3,000bp. CHEER achieves the best performance on identifying viral sequences longer than 1,000bp except the precision for length between 1,000bp and 3,000bp. For short sequences (< 1,000bp), DETIRE has a superior performance. For all lengths, DETIRE achieves the highest accuracy, recall and F1 score than the other three methods.

### 2.6 Performance on the real human gut metagenome

The accuracies, recalls, precisions and F1 scores of DETIRE, CHEER, PPR-Meta, and DeepVirFinder are calculated according to the number of correctly and incorrectly identification viral and host sequences from the real human gut metagenome dataset (Table **??**). DETIRE also achieves a better performance than the other three methods on identifying short viral sequences (<500bp). In spite of a 0.0013 lower precision than CHEER for length between 500bp and 1,000bp, DETIRE gets 0.0043, 0.0117 and 0.0052 higher accuracy, recall and F1 score. For sequences longer than 1,000bp, CHEER is the best performing method. For all lengths, DETIRE achieves higher accuracy, recall and F1 score than PPR-Meta, and DeepVirFinder. Since DETIRE identified less viral sequences longer than 1,000bp than these shorter than 1,000bp, the overall accuracy, recall and F1 score of DETIRE are lower than CHEER.

### 2.7 Comparison on the testing time consuming

The testing time of the four methods on the testing dataset, the CAMI Marine metagenome and the real human gut metagenome are made a comparison in Table **??**. The equipment used for the analysis is two Intel Xeon Gold 6226R (CPU) with the memory of 256Gb. For all of the three datasets, DETIRE has the minimum time consumption for the testing strategies.

### 2.8 Effectiveness of the GCN-based sequence embedding method and the hybrid deep learning based architecture

The sequences from the same training and testing datasets are one-hot encoded by two separate ways: each base in a sequence is one-hot encoded to [1,0,0,0], [0,1,0,0], [0,0,1,0] or [0,0,0,1]; each 3-mer fragment in a sequence is one-hot encoded to a 64-dimension vector. Then the two one-hot encoded training datasets are used to train two deep learning models, namely BOHEM (base one-hot encoded model) and FOHEM (3-mer fragments one-hot encoded model). The two models both have the same hybrid deep learning model as DETIRE. Another two single deep learning models, containing CNN-path and BiLSTM-path respectively, are trained and tested by the same training and testing datasets. The sequences in the datasets are embedded by the GCN-based sequence embedding method from DETIRE. Every model has the same hyper-parameters as the corresponding deep learning model in DETIRE. All parameters are finetuned during the training strategy.

The four models are tested by the testing datasets and made a comparison with DETIRE, which is shown in Fig 4. The accuracy of DETIRE on identifying viral sequences exceeds that of BOHEM and FOHEM by 3.62% and 4.39%, respectively, representing the effectiveness of the GCN-based sequence embedding method in DETIRE. DETIRE also has a superiority than single CNN-based and BiLSTM-based model. The gap is 2.26% and 1.74% on identifying viral sequences, respectively, showing the availability of the hybrid deep learning model in DETIRE.

**Fig 4.**
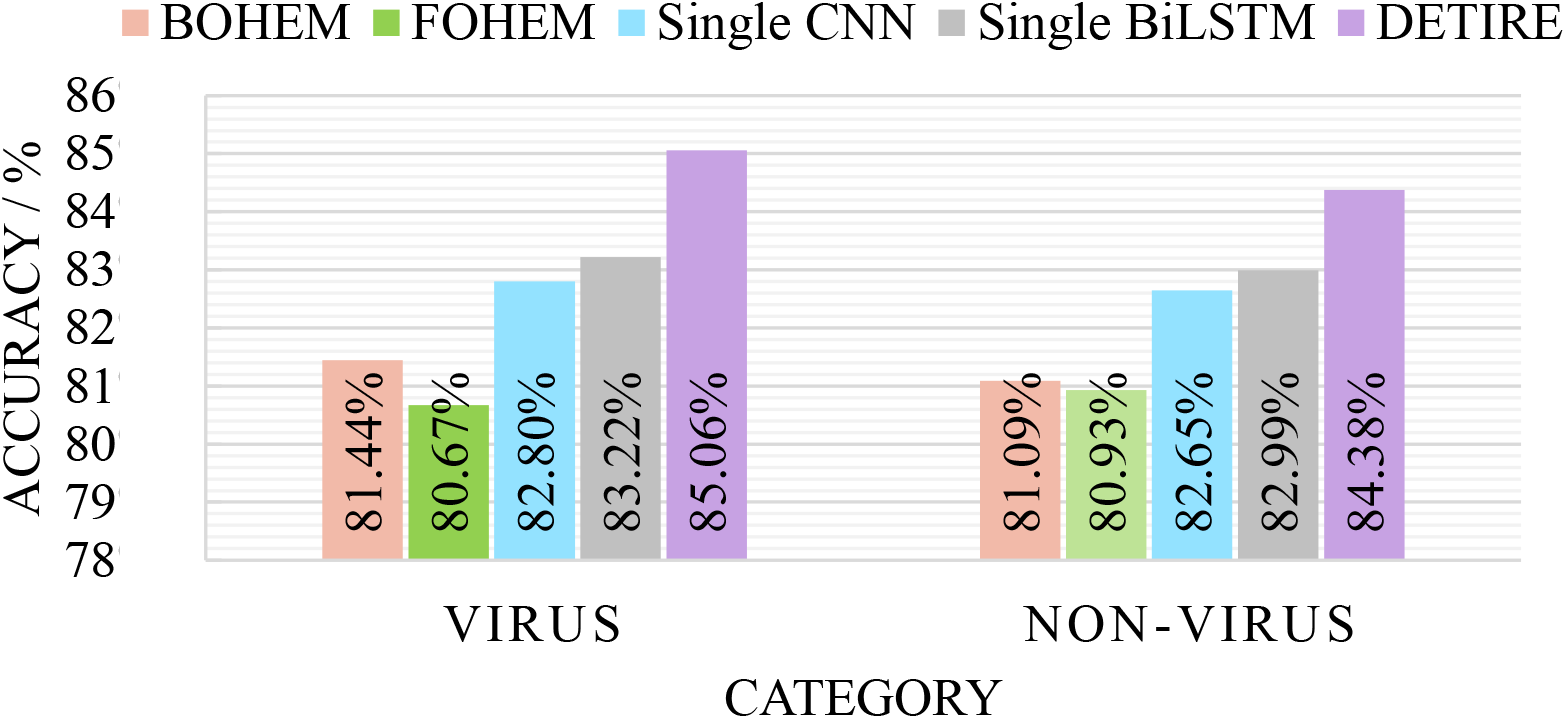
The identification result of BOHEM, FOHEM, single CNN, single BiLSTM and DETIRE on the testing dataset. The five models are tested by the testing dataset to proof the effectiveness of the GCN-based sequence embedding method and the hybrid deep learning based architecture.

## 3 Conclusions

In this paper, a deep learning-based hybrid model, DETIRE, is proposed to identify viral sequences directly from metagenome. Encoded by a graph-based embedding method, nucleotide sequences are fed into a CNN-path and a BiLSTM-path respectively for feature extracting, before being classified by a softmax layer. In comparison to three latest viral identification methods, DeepVirFinder, PPR-Meta and CHEER, on the test dataset, the CAMI Marine dataset and a real human gut metagenome, DETIRE outperforms on identifying short sequences (¡1,000bp). DETIRE will play significant roles in the natural viral community analysis because of the huge number of short sequences generated by the NGS technique. DETIRE is anticipated to play a positive role in the downstream viral analysis such as viral taxonomy and pathogens identification.

## Acknowledgments

This research was funded by the Youth Science and Technology Talent Support Project of Jilin Province grant number QT202109 andthe project funded by China Postdoctoral Science Foundation grant number 2019M651204.

## Notes

### Competing Interest Statement

The authors have declared no competing interest.

